# ZNF512B associates with mitotic spindles, regulates metaphase exit and is crucial for stem cell differentiation

**DOI:** 10.1101/2025.08.21.671475

**Authors:** Lena W. Paasche, Nadine Daus, Johannes J. Groos, Felix Diegmüller, Carlotta Kreienbaum, Erekle Kobakhidze, Jie Lan, Jörg Leers, Stefanie Wunderlich, Annette Borchers, Joel P. Mackay, Tim Marius Wunderlich, Sandra B. Hake

**Affiliations:** Institute for Genetics, Justus-Liebig University Giessen, Germany; School of Life and Environmental Sciences, University of Sydney, New South Wales 2006, Australia; Department of Biology, Molecular Embryology, Marburg University, Germany

**Author notes:** Corresponding authors: Tim Marius Wunderlich and Sandra B. Hake, Institute for Genetics, Justus-Liebig University Giessen, Heinrich-Buff-Ring 58-62, 35392 Giessen, Germany, E-Mails and, phones: 0049 (0)641 99 35473 (TMW) or 35460 (SBH). Shared authorship (Nadine Daus, Johannes J. Groos).

**Keywords:** ZNF512B, mitotic spindle, metaphase exit, stem cell differentiation, NuRD

## Abstract

Zinc finger proteins are a large family of DNA-binding factors that play key roles in diverse cellular processes including gene regulation, RNA metabolism and cell cycle control.

The zinc finger protein ZNF512B has recently been implicated in chromatin organization and transcriptional repression through its direct interaction with the nucleosome remodeling and deacetylase (NuRD) complex, its DNA-binding ability, and its association with the histone variant H2A.Z. Here, we uncover a previously unrecognized role for ZNF512B that is independent of both its zinc finger domains and NuRD association. We identify ZNF512B as a spindle-associated factor that regulates progression through mitosis, specifically controlling metaphase exit. ZNF512B’s N-terminal internal region, which contains 25 repeats of a six-residue motif predicted to form a β-helix structure, is required and sufficient for its spindle interaction. Elevated ZNF512B levels result in a profound metaphase arrest that is ultimately lethal, a phenotype arising from the combined activity of its spindle-binding and chromatin-tethering functions. Conversely, ZNF512B depletion accelerates stem cell proliferation, impairs differentiation, and upregulates genes linked to cell cycle progression.

Our findings position ZNF512B as a multifunctional protein that acts as a transcriptional repressor, a chromatin aggregator and a novel metaphase exit regulator through spindle fiber binding.

## Introduction

Zinc finger (ZF) proteins represent one of the largest protein families in eukaryotes, and are characterized by conserved sequence motifs that fold in small ordered domains stabilized by coordinated zinc ions (1,2). These domains often mediate nucleic acid binding (3–6), enabling many ZF proteins to function as sequence-specific transcription factors that regulate gene expression during development, differentiation, and cell cycle progression (1). Beyond their classical role in transcriptional regulation, a growing number of ZF proteins have been found to exert non-transcriptional functions, including direct involvement in RNA metabolism, chromatin remodeling, protein homeostasis, and even cytoskeletal regulation (1,3,7). Recent studies have demonstrated that certain ZF proteins can also localize to mitotic structures such as the spindle apparatus, where they may contribute to chromosome segregation and mitotic fidelity independent of any DNA-binding activity (8,9). These findings suggest that ZF proteins are functionally versatile regulators involved in both nuclear and cytoplasmic processes essential for maintaining genome stability and cellular homeostasis.

Previously, using label-free quantitative mass spectrometry (lf-qMS) screens we identified ZNF512B to be associated with the evolutionary conserved histone variant H2A.Z (10,11), as well as with H2A.Z’s binding partners PWWP2A (11) and HMG20A (12). ZNF512B, formerly known as GAM/ZFp (13), is a vertebrate-specific ZF protein. It contains eight ZF domains, which are arranged in four pairs that each comprise one atypical C2HC followed by one typical C2H2 sequence. ZNF512B also contains a large internal region (I) with low complexity and a nuclear localization signal (NLS). Only a few studies of ZNF512B function are available so far; these demonstrate a role for the protein in gene regulation and, possibly as a result, in cell homeostasis (13). Further, ZNF512B and its paralogue ZNF512 were found to initiate heterochromatin formation at repetitive elements and pericentric regions (14). It has also been suggested that mutations in regulatory sequences of the *ZNF512B* gene may have a role in the neurodegenerative disease Amyotrophic Lateral Sclerosis (ALS) (15–17).

We recently discovered a conserved amino acid motif within ZNF512B that resembles a consensus sequence known to act as an interaction motif for the nucleosome remodeling and deacetylase (NuRD) complex. Such NuRD-interaction motifs (NIMs) have been previously identified in the N-terminus of the transcription factor Friend of GATA-1 (FOG-1) and several other regulators of gene expression (18–20). We demonstrated that this consensus motif, which is more variable in its composition than previously thought, is both sufficient and required for direct and high-affinity RB Binding Protein 4 (RBBP4) interaction and thereby NuRD complex binding (21). We further identified ZNF512B as a repressor of gene expression that can act in both NuRD-dependent and -independent ways. Surprisingly, high levels of ZNF512B expression lead to chromatin aggregation foci that form independent of ZNF512B’s interaction with the NuRD complex but depend on its ZF domains.

Here we report another unexpected function for ZNF512B – one that is both zinc finger- and NuRD-independent. We identify ZNF512B as a mitotic spindle-associated factor that regulates metaphase exit. An internal proline-rich region containing 25 repeats of a six-residue motif predicted to form a highly basic three-faced β-helix is sufficient for ZNF512B to bind the mitotic spindle. Elevated levels of ZNF512B cause a severe and fatal metaphase arrest, a striking phenotype that primarily depends on ZNF512B’s spindle-binding ability combined with its chromatin tethering function. Conversely, depletion of ZNF512B leads to accelerated stem cell proliferation and impairs the ability of these cells to differentiate into either beating cardiomyocytes (CM) or neuronal progenitor cells (NPCs), as further evidenced by the deregulation of numerous gene expression programs involved in, among other processes, cell cycle control, neurogenesis, and muscle contraction.

In conclusion, our findings reveal that ZNF512B functions not only as a chromatin aggregator and transcriptional repressor but also as a novel regulator of mitosis through its association with spindle fibers. Moreover, ZNF512B plays a critical role in orchestrating stem cell proliferation, migration and differentiation, highlighting its multifunctional role in coordinating cell division and developmental processes.

## Materials & Methods

### Cell Culture and Transfections

HeLa Kyoto (HeLaK) cells were grown in Dulbecco’s modified Eagle’s medium (DMEM, Gibco) supplemented with 10% fetal calf serum (FCS; Gibco) and 1% penicillin/streptomycin (37°C, 5% CO2) and routinely tested for mycoplasma contamination via PCR. Transfections of HeLaK cells were performed using FuGENE® HD Transfection Reagent (Promega) according to the manufacturer’s instructions. Unless stated otherwise, cells were harvested by trypsinization for various assays 48 h after transfection. Transfections of mESCs (v6.5 strain) were performed using jetPEI DNA Transfection Reagent (Polyplus) according to the manufacturer’s protocol. 24 h after transfection, puromycin (0.25 µg/ml) was used to select mESCs for at least 10 days. Finally, mCherry-positive mESC colonies were characterized by genomic PCR, RT-qPCR and immunoblotting, respectively.

Naïve mESCs were cultivated in 2iLIF condition as previously described (12). Specifically, naïve mESCs were cultured on 6-well plates coated with 0.1% gelatine in 2iLIF Medium (N2B27 medium supplemented with 50 μM β-mercaptoethanol, 2 mM L-glutamine, 0.1% sodium bicarbonate, 0.11% bovine serum albumin fraction V, 1000 units/ml recombinant mouse leukemia inhibitory factor (LIF), 1 µM PD0325901 (MEK inhibitor), 3 µM CHIR99021 (GSK3 inhibitor)). Every two days, cells were passaged using Accutase (Stemcell) at a density of 1x10^5^ cells/well.

Embryoid body (EB)-mediated differentiation into cardiomyocytes (CMs) was performed as previously described (12). Briefly, naïve mESCs (Day 0) were primed using differentiation medium (DMEM, Gibco supplemented with 10% FCS, 2 mM L-glutamine, 1% nonessential amino acids, 0.1 mM β-mercaptoethanol, 1000 U/mL LIF) for 2 days. Hanging drops containing 1000 cells in 25 µl of differentiation medium (without LIF, supplemented with 50 µg/ml vitamin C) were prepared and cultured for 4 days. Afterwards, droplets containing EBs were carefully transferred to a 24-well plate coated with 0.1% gelatine and further cultured for 1.5 days in differentiation medium (supplemented with 50 µg/ml vitamin C). Beating cardiomyocytes were observed starting from differentiation Day 7.

Neural progenitor cell (NPC) differentiation was carried out based on the protocol by Bibel et al. (22). Briefly, naïve mESCs were primed as mentioned above. Hanging drops containing 1,000 cells were prepared in 25 µl differentiation medium (without LIF) for 4 days. Then, EBs were pooled and cultured on 100/20 mm bacterial plates in suspension in differentiation medium (without LIF, supplemented with 5 µM retinoic acid (RA)) for 4 days. Medium was changed every two days. On differentiation Day 10, the generated EBs containing NPCs were collected for further analyses.

### Plasmid Cloning

Human ZNF512B full-length and deletion plasmids were obtained from (21). Additional GFP-ZNF512B_IN and _IC plasmids were constructed as described in (21). *Xenopus laevis* Znf512b plasmid was generated via NEBuilder HiFi DNA Assembly (NEB). pc1116-H2B-mRFP (23) was a gift from Prof. Dr. M. Cristina Cardoso (Cell Biology and Epigenetics, Department of Biology, Technical University of Darmstadt, Germany). *Znf512b* KO mESC cell lines were established using CRISPR/Cas9 technology. Briefly, sgRNAs were designed to target the third exon of *Znf512b* using the online tool CRISPOR, synthesized by Integrated DNA Technology (IDT), and cloned into the vector pX461 (Addgene). Donor pUC19-based vectors containing either mCherry or a puromycin resistance and mammalian transcriptional triple terminators bGH + hGH + SV40 (synthesized by GENEWIZ) surrounded by homology arms were constructed via NEBuilder HiFi DNA Assembly (NEB). Homology arms were generated from mESC genomic DNA obtained using the QIAamp DNA Mini Kit (QIAGEN) via PCR using Q5 High-Fidelity DNA Polymerase (NEB). All relevant sgRNA sequences and primers are listed in the **Supplementary Information**.

### Primers and Oligos

All primers and oligos are listed in the **Supplementary Information.**

#### Immunofluorescence (IF) Microscopy

Immunofluorescence (IF) staining of cells and their microscopic analysis was performed as previously described (12). Briefly, 1x10^5^ adherent HeLaK cells expressing GFP, GFP-ZNF512B or its deletions/mutants were seeded on coverslips in 24-well plates and cultured overnight (ON). The next day, cells were washed two times with PBS and fixed for 15 min in PBS containing 1% (v/v) formaldehyde. After washing, cells were permeabilized and blocked with PBS containing 0.1% (v/v) Triton™ X-100 (PBS-T) and 1% (w/v) bovine serum albumin (BSA) for 20 min. Cells were incubated stepwise with primary and then secondary antibodies in PBS-T + 1% BSA for 30 min with three wash-steps using PBS-T in between. After three final PBS-T wash-steps, DNA was stained with 10 µg/ml Hoechst solution for 3 min. After washing with H2O, coverslips with cells were then mounted in Fluoromount-G® mounting medium (SouthernBiotech) on microscope slides (Epredia). Images were acquired using an Axio Observer.Z1 inverted microscope (Carl Zeiss) with an Axiocam 506 mono camera system. Image processing was performed with Zeiss Zen (version 3.1, blue.edition) and ImageJ2 (version 2.14.0/1.54f).

Live-cell imaging to monitor mitotic progression of HeLaK cells expressing RFP-H2B and various GFP-ZNF512B constructs was executed with a CELLCYTE X (Cytena) microscope using a 10x objective. Cells were transfected as described above. Imaging started 4-6 h post-transfection with the following set up: 16 images per well were taken every 20 min for 36 h with 150 ms exposure time for the green channel (Excitation: 473-492 nm, Emission: 502-561) and 450 ms exposure time for the red channel (Excitation: 580-598, Emission: 612-680 nm). Image processing was performed with Zeiss Zen (version 3.1, blue.edition), ImageJ2 (version 2.14.0/1.54f) and Imaris (version 9.9.0). For more detailed 3D visualization, a Zeiss Spinning-Disc microscope system (Axio Observer.Z1) equipped with a 100x objective was used.

For growth curve analyses, cells were imaged every 2 h for 4 days. Live-cell imaging movies and analyses of growth curves were generated with the Cytena software.

#### Antibodies

All antibodies are listed in the **Supplementary Information.**

#### DNase I-Immunoprecipitation (DNase I-IP) after Nocodazole Treatment

HeLaK cells were seeded one day before transfection on 145/20mm plates (Sarstedt). 24 h after transfection, cells were treated with 100 ng/ml nocodazole for 22 h. Cells were harvested by collecting supernatant and by washing remaining mitotic cells off from plate with 1 x PBS. Treatment efficiency was analyzed with propidium iodide (PI) staining followed by analysis with a flow cytometer (BD Accuri C6 plus). For PI staining, cells were fixed with 70% methanol and stored at -20°C for at least 1 h up to several days. After washing, cells were incubated in PI staining solution (100 µg/ml RNaseA, 50 µg/ml PI) for 30 min at 37°C in dark and analyzed.

Immunoprecipitation was performed using GFP-Trap® magnetic particles M-270 (Chromotek) following the manufacturer’s protocol. In brief, cells were incubated for 30 min on ice with 200 µl RIPA buffer (10 mM Tris/Cl pH 7.5, 150 mM NaCl, 0.5 mM EDTA, 0.1% SDS, 1% Triton™ X-100, 1% deoxycholate) supplemented with DNase I (75 Kunitz U/mL, Thermo Fisher Scientific) and 2.5 mM MgCl2. After centrifugation at 17,000 g for 10 min at 4°C, lysate was transferred to a fresh tube and mixed with 300 µl dilution buffer (10 mM Tris/Cl pH 7.5, 150 mM NaCl, 0.5 mM EDTA). Lysate was then incubated with GFP-Trap® magnetic particles M-270 (Chromotek) rotating for 2 h at 4°C. After incubation, beads were washed three times with wash buffer (10 mM Tris/Cl pH 7.5, 150 mM NaCl, 0.05% Nonidet^TM^ P-40 Substitute, 0.5 mM EDTA), and precipitated proteins were eluted by boiling beads for 5 min in 2x SDS-sample buffer. Eluates were compared to input material via semi-dry immunoblotting.

#### RNA Extraction, RT-qPCR and RNA-seq

RNA extraction was performed as described (24). Briefly, total RNA was extracted using the RNeasy Mini Kit (QIAGEN) with on column DNase digest (RNase-Free DNase Set, QIAGEN) according to the manufacturer’s instructions. 1 µg total RNA was reverse transcribed using the Transcriptor First Strand cDNA Synthesis Kit (Roche) with random hexamer primers according to the manufacturer’s protocol. For subsequent qPCR analysis, technical triplicates of 3 µl cDNA (1:20 dilution), 7.5 µl iTaq™ Universal SYBR® Green Supermix (Bio-Rad), 3 µl water and 1.5 µl primer mix (5 µM forward and reverse primer) were used. The qPCR program consisted of 5 min at 95^°^C, followed by 40 cycles at 95^°^C for 3 sec and 60^°^C for 20 sec. Lastly, Ct values of technical triplicates were averaged, and fold change expression was calculated with the Delta-Delta Ct method, normalizing to control samples and HPRT1 expression. mRNA sequencing (mRNA-seq) was performed at Biomarker Technologies (Germany).

#### Structure Predictions

Protein structures were predicted using either AlphaFold2 implemented as ColabFold (https://colab.research.google.com/github/sokrypton/ColabFold/blob/main/AlphaFold2.ipynb) or the AlphaFold3 server (https://alphafoldserver.com/) and visualized in Pymol 3.1 (Schroedinger software).

#### mRNA-seq Analysis

mRNA-seq analysis was performed on the Galaxy platform (25). Raw sequencing files (FASTQ) were adapter- and quality-trimmed using trimGalore (26). Alignment of the trimmed sequencing reads to the mm9 reference genome (Illumina’s iGenomes) was performed using Hisat2 v.2.2 (27). Read counts per gene were obtained from BAM files using featureCounts (28) from the Rsubread package (29) and the mouse mm9 gene transfer format (GTF).

Normalization of read counts and detection of differentially expressed genes (DEGs) was performed with DESeq2 (30) v1.46.0 in R v4.4.1 (https://www.R-project.org/). If not otherwise indicated, significant DEGs were chosen based on an adjusted p-value <0.05 (**Supplementary Tables 1 and 2**). Based on the resulting DEGs, Venn diagrams were generated using the eulerr (https://CRAN.R-project.org/package=eulerr) and ggplot2 (https://ggplot2.tidyverse.org) packages. The overlap between KO clones and cell stages was used for an over-representation gene ontology (GO) (31,32) and KEGG pathway (33–35) analysis with clusterProfiler (36). Volcano plots were generated with VolcanoPlot on the galaxy server (37).

## Results

### ZNF512B localizes to mitotic spindles via its internal region and independent of NuRD interaction

We recently reported that overexpression of the zinc finger protein ZNF512B, fused either N- or C-terminally to a GFP tag (GFP-ZNF512B or ZNF512B-GFP), leads to the formation of distinct nuclear foci, the size of which correlates with ZNF512B expression levels (21). This chromatin aggregation phenotype is driven by a combination of ZNF512B’s zinc finger-dependent abilities to oligomerize and to bind DNA. Upon closer examination of GFP-ZNF512B expressing cells outside of interphase, we observed a striking shift in its subcellular localization; a large portion of the GFP-ZNF512B protein pool re-localized from chromatin to the mitotic spindle in metaphase, with only a minor fraction remaining associated with condensed chromosomes (**Figure 1A, Supplementary Figure S1A**). This spindle localization was further confirmed in greater detail by live-cell imaging of GFP-ZNF512B in cells transiently expressing red fluorescent protein (RFP)-tagged histone H2B to visualize chromatin (**Movie 1**).

**Figure 1:**
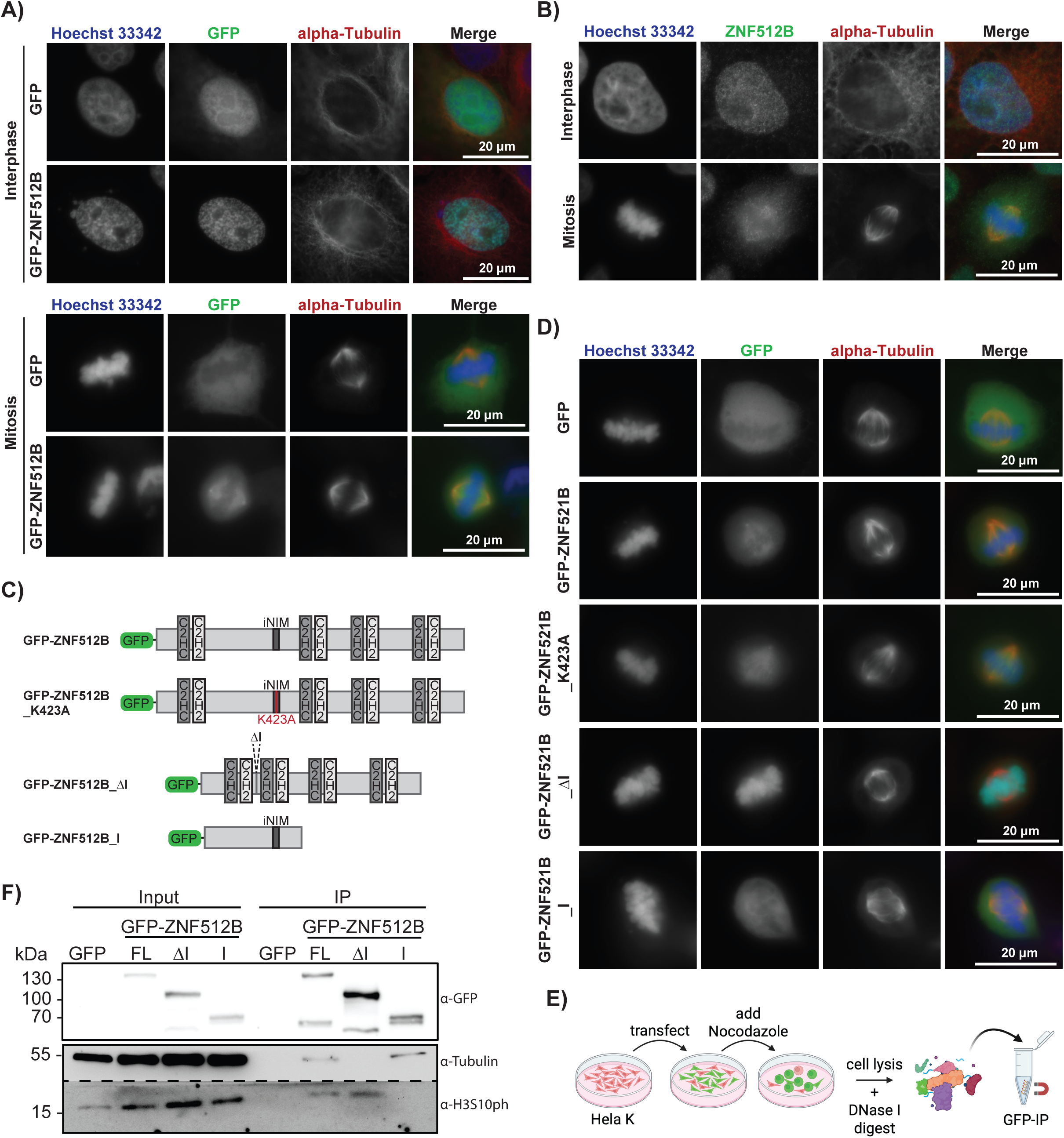
ZNF512B associates with mitotic spindles via its internal region, independently of NuRD binding. **(A)** Immunofluorescence microscopy pictures of HeLaK cells during interphase (top) and metaphase (bottom) transfected with GFP (control) or GFP-ZNF512B (green). DNA (blue) was visualized by Hoechst 3342 and microtubules/mitotic spindles by anti-alpha-tubulin (red) staining. Scale bar = 20 µm. See also **Movie 1**. **(B)** Immunofluorescence microscopy pictures of HeLaK cells stained with Hoechst 3342 (DNA, blue), anti-ZNF512B (green) and anti-alpha-tubulin (microtubules/mitotic spindle; red). Scale bar = 20 µm. **(C)** Schematic depiction of GFP-ZNF512B, its lysine 423 to alanine substitution mutant (K423A) and various deletion constructs. **(D)** Immunofluorescence microscopy pictures of HeLaK cells during interphase (top) and metaphase (bottom) transfected with GFP (control), GFP-ZNF512B WT or point mutant and deletion constructs (green). DNA (blue) was visualized by Hoechst 3342 and microtubules/mitotic spindles by anti-alpha-tubulin (red) staining. Scale bar = 20 µm. **(E)** Schematic depiction of experimental layout to identify mitosis-specific binding partners of GFP-ZNF512B. **(F)** Immunoblots using extracts generated through DNase I digestion from Nocodazole treated HeLaK cell transfected with GFP-ZNF512B constructs (see Figure 1E) using antibodies against GFP, alpha-tubulin (spindle component) and histone H3 serine 10 phosphorylation (H3S10ph; PTM of mitotic chromatin). FL: Full Length.

Given the strong overexpression of GFP-ZNF512B protein in our experimental system (approximately 500-fold enrichment of ZNF512B mRNA compared to endogenous levels (21)), we next asked whether the observed mitotic spindle localization might represent an overexpression artifact. To address this question, we performed immunofluorescence (IF) microscopy with an antibody against endogenous ZNF512B (21). Consistent with the localization of overexpressed GFP-ZNF512B, endogenous ZNF512B protein was also clearly detected at the mitotic spindle apparatus in metaphase, anaphase and at the midbody in cytokinesis (**Figure 1B, Supplementary Figure S1B**). This finding confirms that ZNF512B’s association with spindle fibers during mitosis is not an experimental artifact.

Next, we investigated which region in ZNF512B is required for spindle binding by analyzing the mitotic localization of several GFP-ZNF512B mutant constructs that we had previously functionally described (21) (**Figure 1C**). A key focus was to assess whether spindle recruitment requires ZNF512B’s direct interaction with the NuRD complex, which is mediated by its internal variant NuRD interaction motif (iNIM) (21). Several components of NuRD, such as CHD4 (38) and MTA1 (39), have been reported to localize to the mitotic spindle, and the NuRD subunit RBBP4 (to which ZNBF512B directly binds) has been shown to be essential for bipolar spindle assembly during meiotic metaphase (40).

To evaluate the potential role of the NuRD complex in recruiting ZNF512B to the mitotic spindle, we used the K423A point mutant of ZNF512B (**Figure 1C**), which is unable to bind NuRD (21). We also examined GFP-ZNF512B constructs with either all eight zinc fingers deleted – retaining only the internal region (I) – or with the internal region containing the iNIM deleted – retaining all zinc fingers (ΔI) (**Figure 1C**). Unexpectedly, both GFP-ZNF512B_K423A and GFP-ZNF512B_I mutants retained spindle-binding ability (**Figure 1D**). In contrast, GFP-ZNF512B_ΔI failed to associate with the spindle apparatus (**Figure 1D**), indicating that the internal region of ZNF512B is both necessary and sufficient for mitotic spindle localization, and that this interaction is NuRD independent.

This surprising finding was further supported in immunoprecipitation (IP) experiments using extracts from DNase I-treated cells arrested in metaphase by Nocodazole, a microtubule polymerization inhibitor (**Figure 1E**). In line with our previous findings, GFP-ZNF512B co-precipitated both mitotic chromatin (histone H3 serine 10 phosphorylation; H3S10ph) and alpha-tubulin, one major spindle component (**Figure 1F**). The zinc finger regions of ZNF512B, lacking the internal domain, precipitated only chromatin but not spindle-associated tubulin. Conversely, ZNF512B’s internal domain alone was sufficient to bind spindle components but did not associate with chromatin.

Next, we investigated whether ZNF512B is responsible for recruiting NuRD complex members to the mitotic spindle. Overexpression of GFP-ZNF512B_I and GFP-ZNF512B_ΔI constructs revealed that the relatively weak localization of endogenous CHD4 to spindle fibers remained, regardless of the ZNF512B’s ability to bind the NuRD complex (**Supplementary Figure S1C**). Interestingly, ZNF512B’s direct binding partner RBBP4 showed increased localization to spindle fibers when ZNF512B’s internal region containing the iNIM was present, compared to the ΔI construct, which is unable to associate with either the mitotic spindle or RBBP4 (**Supplementary Figure S1D**). These results suggest that ZNF512B may not be required for the recruitment of all NuRD complex components to the spindle apparatus, but does play a role in the spindle association of its direct interaction partner, RBBP4.

In summary, we have identified the internal region of ZNF512B to be necessary and sufficient for its association with mitotic spindles, independently of NuRD complex interaction.

### ZNF512B’s N-terminal internal region is sufficient for spindle interaction

To further refine the region of ZNF512B responsible for spindle binding, we first examined the evolutionary conservation of ZNF512B’s internal region, which comprises a low-complexity sequence (21). Interestingly, only a C-terminal portion of the internal region (IC) is conserved across all vertebrates, whereas an N-terminal portion (IN) is specific to mammals (**Supplementary Figure S2**). Visual inspection of the sequence of ZNF512B’s IN region revealed 25 tandem copies of a six-residue motif that broadly follows the consensus sequence [[KR]-P-φ-[Pπ]-φ-π (where φ is an aliphatic hydrophobic residue and π is a small, polar residue; **Figure 2A**). This region was conserved overall in other mammalian orthologues, though with the insertion or deletion of one or more repeat motifs compared to the human protein. Prediction of the structure of the human IN region using AlphaFold2 suggests, albeit with moderate confidence, that this region forms an unusual three-faced, right-handed β-helix (**Figure 2B, Supplementary Figure 3A**).

**Figure 2:**
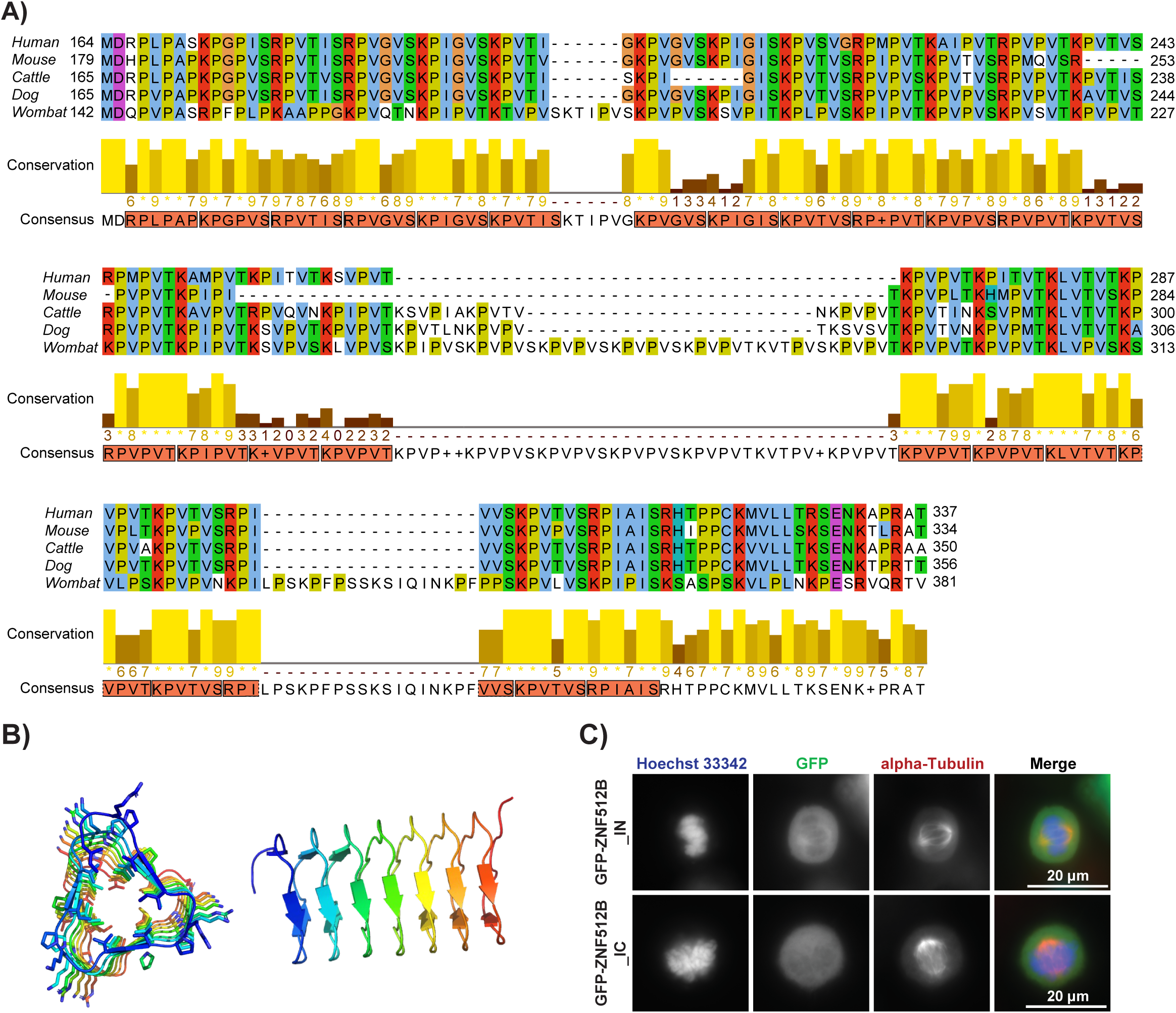
The mammalian-specific N-terminal portion of ZNF512B’s internal region forms a beta-helix structure and is sufficient and required for spindle association. **(A)** Amino acid alignment of the N-terminal portion of ZNF512B’s internal region across mammalian and marsupial (Wombat) species with the SnapGene (Version 8.0.3) Software using Clustal-Omega and Jalview (Version 2.11.4.1). Protein Sequences were obtained from uniprot.org for human (Homo sapiens, Entry Q96KM6), mouse (Mus musculus, Entry Q6ZPW1), cattle (Bos taurus, Entry G3N139), dog (Canis lupus familiaris, Entry A0A8P0NRY1) and wombat (Vombatus ursinus, Entry A0A4X2L7X1). Consensus sequence and conservation score were generated using Jalview Version 2.11.4.1. Amino acids are colored in the Clustal X default coloring scheme: hydrophobic (blue), positive charge (red), negative charge (magenta), polar (green), cysteine (pink), glycine (orange), proline (yellow), aromatic (cyan) and unconserved or gap (white). Repeat regions are marked by oranges boxes in the consensus sequence. **(B)** AlphaFold2 prediction of the folding of ZNF512B’s IN region. Shown are two views rotated 90°. **(C)** Immunofluorescence microscopy pictures of HeLaK cells during metaphase transfected with GFP-ZNF512B_IN (IN) or GFP-ZNF512B_IC (IC) constructs (green). DNA (blue) was visualized by Hoechst 3342 and microtubules/mitotic spindles by anti-alpha-tubulin (red) staining. Scale bar = 20 µm.

Based on these sequence alignments, we generated GFP-tagged constructs containing either the IN or IC region of ZNF512B; with the IN region fused to a nuclear localization signal (NLS) sequence to ensure nuclear localization of both constructs as the endogenous NLS is predicted to be located solely within the IC domain (**Supplementary Figure S2**). Surprisingly, upon transient transfection, only the mammalian-specific IN but not the evolutionary conserved IC region was able to associate with spindle fibers during metaphase (**Figure 2C**), indicating that IN is sufficient for spindle interaction.

In line with this finding, GFP-tagged *Xenopus laevis* Znf512b protein, which naturally lacks the IN domain (**Supplementary Figure S2**), did not localize to the mitotic spindle nor was it found in the midbody, although it was still able to bind condensed chromatin during mitosis and to form ZF-dependent chromatin aggregates in interphase, when expressed in human HeLaK cells (**Supplementary Figure S3A**).

In conclusion, we identified a mammalian-specific N-terminal internal region of ZNF512B, which may adopt a β-helix structure, as both necessary and sufficient for its interaction with the mitotic spindle.

### Overexpression of GFP-ZNF512B results in mitotic arrest dependent on IN-mediated spindle binding

Next, we investigated whether elevated levels of ZNF512B and/or its spindle-binding ability affect mitotic progression. We transfected either GFP alone (control) or diverse GFP-ZNF512B mutant constructs into HeLaK cells stably expressing RFP-tagged H2B and performed live-cell imaging. Cells were monitored over a 36 h time period, with images acquired every 20 min using the CELLCYTE X (Cytena) system. The duration of mitosis was subsequently quantified. Typically, HeLaK cells spend approximately one hour in mitosis, as we also observed in cells expressing GFP alone (**Figure 3A-C, Movie 2**). In contrast, GFP-ZNF512B overexpression caused a severe mitotic arrest lasting up to 25 h, with an average mitotic duration of 17 h, after which most of the cells died (**Figure 3A-C, Movie 3**). We next tested cells expressing the GFP-ZNF512B_K423A point mutant, which cannot bind NuRD but retains the ability to associate with the mitotic spindle. These cells exhibited a similarly pronounced mitotic arrest followed by cell death, indicating that this phenotype is independent of NuRD binding (**Figure 3A-C, Movie 4**). On the other hand, deletion of the spindle-binding internal domain (GFP-ΔI) did not cause any mitotic defects (**Figure 3A-C, Movie 5**), while expression of only the internal domain alone (GFP-I) again led to a mitotic delay, though less severe than observed with the full-length protein (**Figure 3A-C, Movie 6**). In this case, cells were able to survive a delayed mitotic exit, suggesting that excess of ZNF512B and its enhanced spindle association and zinc finger-mediated chromatin binding are particularly detrimental. As expected, overexpression of spindle-binding GFP-IN (**Figure 3A-C, Movie 7**), but not GFP-IC (**Figure 3A-C, Movie 8**), also led to a mild mitotic delay of approximately 2.5 h, further supporting the idea that docking of even a minimal ZNF512B fragment with the mitotic spindle apparatus is sufficient to impair timely mitosis exit.

**Figure 3:**
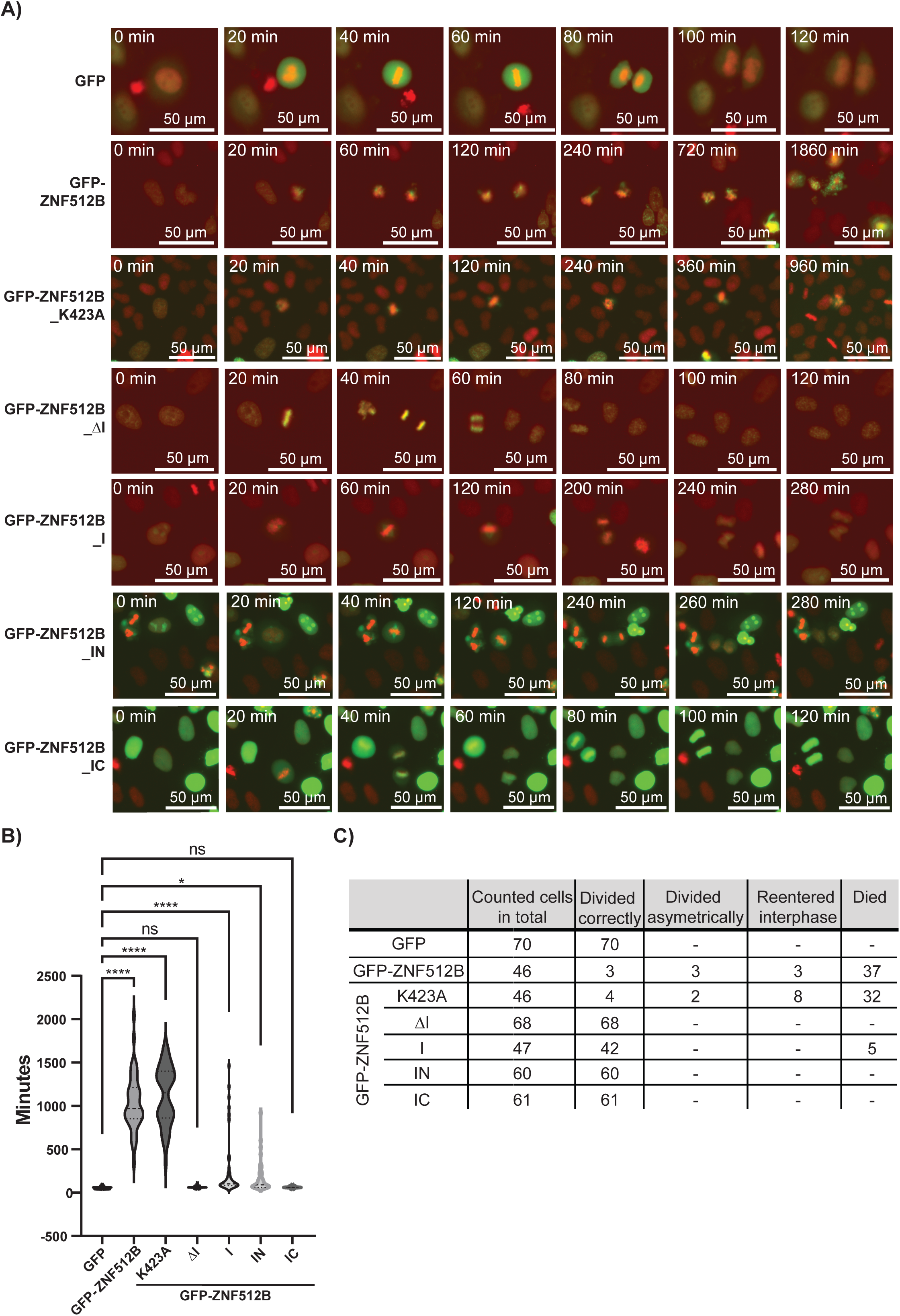
Elevated levels of ZNF512B lead to severe mitotic delay, depending on its IN region. **(A)** Pictures taken at different time points from live-cell imaging movies from HeLaK cells stably expressing RFP-H2B (red) transfected with various GFP-ZNF512B mutant and deletion constructs (green, see Figure 1C). See **Movies 2-8**. Scale bar = 50 µm. **(B)** Quantification of time transfected HeLaK cells (see **A**) spent in mitosis. Significance was calculated in Prism with one-way ANOVA, where ns = > 0.05, * = <0.05, ** = <0.01, *** = <0.001 and ****<0.0001. **(C)** Table summarizing observed phenotypes in transfected cells (see **A**).

Together, these data suggest that high levels of ZNF512B protein disrupt normal mitotic progression, likely through its combined association with the mitotic spindle and chromatin; interactions that are mediated by the N-terminal internal region and the zinc finger domains, respectively.

### ZNF512B depletion leads to a growth advantage and upregulation of genes involved in cell cycle control

Having shown that ZNF512B binds to the mitotic spindle via its N-terminal internal region and that high levels of ZNF512B cause a severe mitotic delay followed by cell death, we next investigated whether depletion of ZNF512B would have the opposite effect on cell division and proliferation. We generated ZNF512B-depleted mouse embryonic stem cells (*Znf512b* KO mESCs) by inserting a triple transcriptional terminator sequence in between the third and fifth exon of the murine *Znf512b* (also termed *Zfp512b*) gene, thereby deleting the fourth exon (**Supplementary Figure S4A**). We obtained six independent *Znf512b* KO mESC clones, confirming their identity at the genomic, RNA and protein levels (**Supplementary Figure S4B-D**). Interestingly, all naïve *Znf512b* KO mESC clones exhibited enhanced proliferation compared to wild type (WT) cells (**Figure 4A**). This striking finding aligns well with our observation that elevated ZNF512B levels cause a mitotic delay, further supporting a role for ZNF512B in the regulation of mitosis in both human and mouse cells.

**Figure 4:**
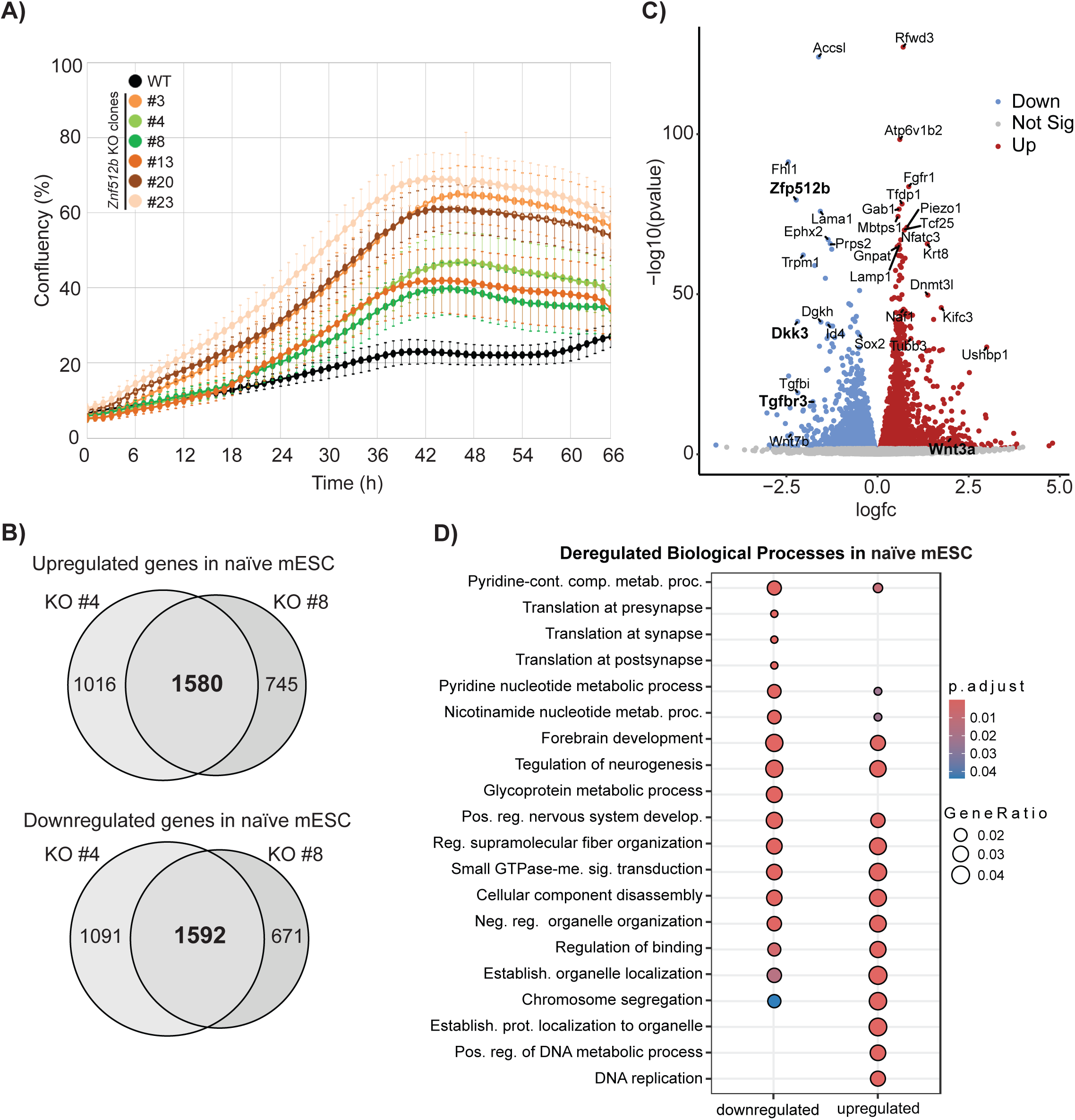
ZNF512B depletion leads to faster proliferating stem cells. **(A)** Growth curve analysis of WT and six naïve Znf512b KO mESC cell clones. **(B)** Venn diagrams of differentially up- or downregulated genes in naïve Znf512b KO mESC clones #4 and #8 compared to naïve WT mESCs. **(C)** Volcano plot of combined deregulated genes in naïve Znf512b KO mESC clones #4 and #8 compared to naïve WT mESCs. **(D)** GO term analysis (‘biological processes’) of combined down- and upregulated genes in both naïve Znf512b KO mESC clones compared to naïve WT mESCs.

To gain insight into the molecular changes underlying these proliferative differences, we performed RNA-sequencing (RNA-seq) of WT mESCs and two *Znf512b* KO clones (#4 and #8) in the naïve state (**Supplementary Figure S4E, Supplementary Table 1**). These two cell clones, which exhibited the least increased proliferation phenotype, were selected to minimize confounding effects arising from potential off-target genomic changes during selection. RNA-seq analysis revealed more than 3,000 genes that were commonly deregulated in both naïve *Znf512b* KO clones compared to WT mESCs, with approximately half being upregulated and the other half being downregulated (**Figure 4B, C**). Using all six previously generated naïve *Znf512b* KO cell clones, these findings were further validated by RT-qPCR for four consistently deregulated genes (**Supplementary Figure S4F**), which are involved in the regulation of cell growth and/or differentiation – *Wnt3a* (Wnt family member 3a; upregulated), *Tgfbr3* (transforming growth factor beta receptor type 3; downregulated), *Dkk3* (Dickkopf WNT signaling pathway inhibitor 3; downregulated) and *Ctnnb1* (Catenin Beta 1; beta-catenin signaling; slightly downregulated).

Gene ontology (GO) analysis of these commonly deregulated genes revealed that downregulated genes were associated with neurogenesis, while both down- and upregulated genes were linked to metabolism and organelle organization (**Figure 4D**). Interestingly, and consistent with our observation that naive *Znf512b* KO mESCs proliferate faster than WT, upregulated genes were associated with cell cycle regulation via the term ‘DNA replication’ (**Figure 4D**).

Together, these data suggest that reduced ZNF512B levels increase cell proliferation and upregulate expression of genes involved in cell cycle progression in naïve mESCs.

### Loss of ZNF512B severely affects cell differentiation and proliferation

Given that naïve *Znf512b* KO mESCs exhibited increased proliferation, we next investigated whether ZNF512B is required for proper mESC differentiation. Using established protocols (12,22), we differentiated WT mESCs and the two *Znf512b* KO cell clones #4 and #8 into either beating cardiomyocytes (CMs) or neural progenitor cells (NPCs) (**Figure 5A**). First, we observed that Day 4 embryoid bodies (EBs) derived from both *Znf512b* KO cell clones were significantly larger than those from WT cells (**Figure 5B, C**). Furthermore, ZNF512B-deficient cells failed to properly differentiate into beating CMs (**Figure 5D**). Differentiation into NPCs resulted in enlarged cell clusters (**Figure 5E**), demonstrating increased proliferation and/or enhanced cell migration (**Figure 5F**).

**Figure 5:**
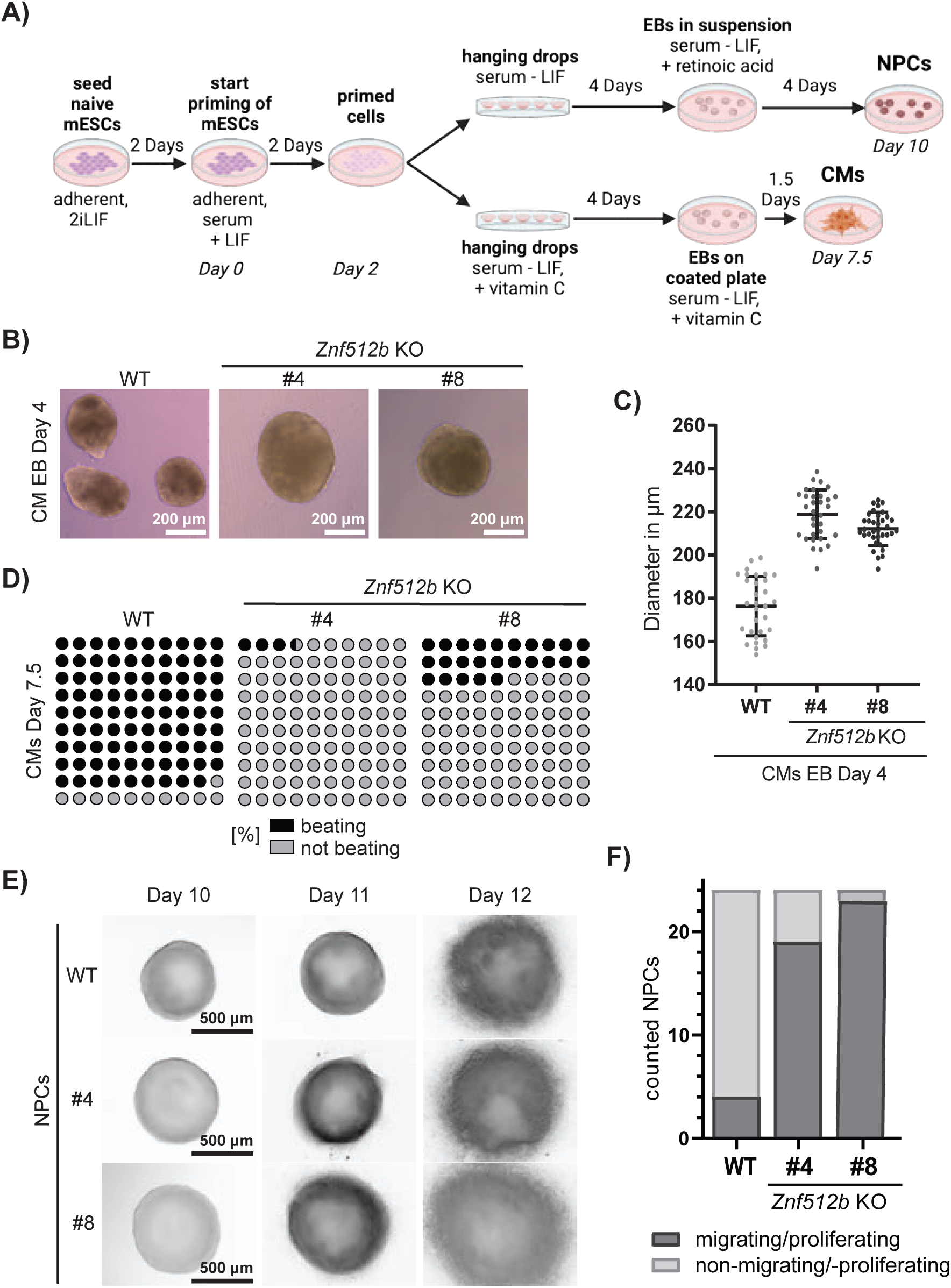
ZNF512B depletion results in differentiation and migration defects. **(A)** Schematic depiction of applied protocols to differentiate mESC into neuronal progenitor cells (NPCs, top) or beating cardiomyocytes (CMs, bottom). **(B)** Microscopic images of Day 4 Embryoid Bodies (EB) during the differentiation process towards CMs derived from WT and Znf512b KO clones #4 and #8. Scale bar = 200 µm. **(C)** Quantification of CM-differentiation protocol derived Day 4 Embryoid Body (EB) sizes of WT and Znf512b KO clones #4 and #8 (see **B**). 31 WT, 32 #4 and 34 #8 EBs were analyzed and diameters of single EBs depicted in µm. Significance was calculated in Prism with one-way ANOVA, where ns = > 0.05, * = <0.05, ** = <0.01, *** = <0.001 and ****<0.0001. **(D)** Depiction of percent beating (black) or non-beating (gray) WT or two Znf512b KO clones #4 or #8 EBs at Day 7.5 of the CM differentiation procedure (see **A**). **(E)** Microscopic images of Day 10, 11 and 12 NPCs derived from WT and Znf512b KO clones #4 and #8. Scale bar = 500 µm. **(F)** Quantification of migrating/proliferating (dark gray) or non-migrating/-proliferating (light gray) NPCs one day after plating Day 10 EBs of WT and two Znf512b KO clones #4 or #8.

RNA-seq analysis of Day 2 primed cells, as well as differentiated Day 7.5 CMs or Day 10 NPCs (**Supplementary Figure S5A, Supplementary Table 2**), identified 203 deregulated genes shared between both *Znf512b* KO clones compared to WT cells across all three differentiation stages (**Supplementary Figure S5B**). The vast majority of these genes was upregulated, with only three genes downregulated (**Supplementary Figure S5C**), supporting the notion, that ZNF512B primarily functions as a transcriptional repressor as previously proposed (14,21). Kyoto Encyclopedia of Genes and Genomes (KEGG) pathway analysis revealed that commonly upregulated genes were predominantly associated with RNA degradation and, notably, cell cycle regulation (**Supplementary Figure S5D**). These findings indicate, once again, that *Znf512b* KO cells display aberrant cell cycle progression throughout all stages of differentiation.

Next, we examined gene expression changes at each individual stage of differentiation. In both primed *Znf512b* KO clones, nearly 1,000 genes were differentially expressed compared to primed WT cells (**Figure 6A**), with approximately two-thirds being upregulated and one-third being downregulated (**Supplementary Figure S6A**). GO analysis revealed that downregulated genes were associated with various metabolic processes, whereas upregulated genes were once again related to pathways involved in cell cycle, such as DNA replication, among others (**Figure 6B**). A quantitatively similar pattern was observed in Day 10 NPCs (**Figure 6C, Supplementary Figure S6B**), although in this context, the affected genes in *Znf512b* KO cells were, as expected, primarily linked to neurogenesis (**Figure 6D**). Notably, genes involved in the Wnt signaling pathway were also deregulated, corroborating our findings in naïve cells (see **Supplemental Figure S4F**) and indicating that this key developmental pathway is disrupted early in development and remains affected during NPC differentiation. The greatest number of commonly affected genes in both Znf512b KO clones was detected in Day 7.5 CMs (**Figure 6E**), with 750 genes being upregulated and 971 being downregulated (**Supplementary Figure S6C**). GO analyses showed that downregulated genes were significantly enriched in terms related to muscle contraction/differentiation and cardiac muscle development (**Figure 6F**), consistent with our finding that loss of ZNF512B impairs the differentiation of functional, beating CMs (see **Figure 5D**). Interestingly, the upregulated genes were again predominantly associated with cell cycle processes, including mitotic nuclear division and sister chromatin segregation (**Figure 6F**).

**Figure 6:**
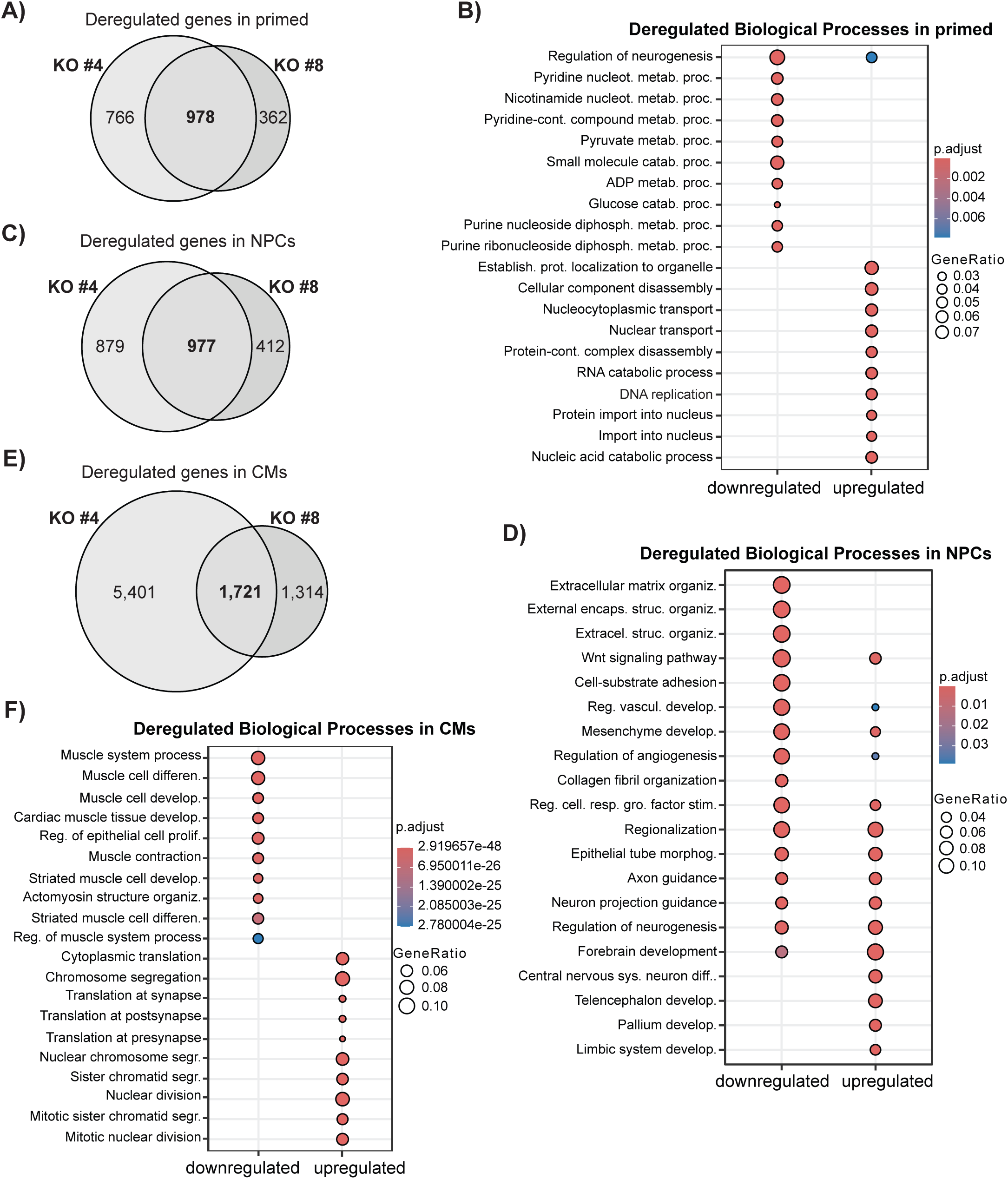
Loss of ZNF512B leads to deregulated gene expression during distinct differentiation pathways. **(A, C, E)** Venn diagrams of differentially expressed genes in Znf512b KO clones #4 and #8 compared to WT cells in the primed (**A**), NPC (**C**) or CM (**E**) stages. **(B, D, F)** GO term analysis (‘biological processes’) of shared deregulated genes in both Znf512b KO clones in the primed (**B**), NPC (**D**) or CM (**F**) stages, corresponding to **A**, **C**, and **E**.

Collectively, these findings underscore the essential role of ZNF512B in regulating cell proliferation, and in ensuring proper stem cell differentiation.

## Discussion

We demonstrate here that ZNF512B binds to mitotic spindle fibers independently of its interaction with the NuRD complex and with DNA/chromatin. This association with main components(s) of the mitotic spindle apparatus is mediated by a mammalian-specific N-terminal internal region that contains a repeat motif predicted to form a β-helix structure. Elevated ZNF512B protein levels result in a pronounced delay in metaphase exit, ultimately leading to cell death. This severe phenotype depends on ZNF512B’s ability to bind both mitotic spindles and chromatin. In contrast, ZNF512B depletion accelerates stem cell proliferation, impairs stem cell differentiation, and induces the upregulation of genes involved in cell cycle control.

ZNF512B’s ability to bind the mitotic spindle does not depend on its interaction with the NuRD complex, although several NuRD components, such as CHD3 (41), CHD4 (38), MTA1 (39) and RBBP4 (40), have been reported to also localize to the spindle apparatus or to cause spindle disorganization when depleted. Moreover, ZNF512B is likely not required for the recruitment of NuRD members to the spindle fiber, as the amount of CHD4 localized to the metaphase spindle is not affected by overexpression of ZNF512B constructs regardless of their ability to bind the NuRD complex. The observed increase in RBBP4 at the mitotic spindle upon overexpression of the I, but not the ΔI, construct may be explained by the fact that RBBP4 is the direct binding partner of ZNF512B via its iNIM. One can speculate that I-dependent higher levels of RBBP4 compared to CHD4 at the spindle may reflect differences in protein expression levels. It is possible that all endogenous CHD4 protein is already bound at the spindle via endogenous ZNF512B, whereas excess of RBBP4 protein, which is normally not associated with the spindle, is additionally recruited through high levels of overexpressed GFP-ZNF512B_I. While these points require further clarification, it is tempting to speculate that most NuRD complex members interact with the mitotic spindle via distinct binding partners. It is therefore likely that they all fulfill separate, non-redundant functions during mitosis.

Spindle binding of ZNF512B is mediated by its mammalian-specific N-terminal internal domain, as demonstrated by experiments with *X. laevis* GFP-ZNF512B. The IN region contains up to 25 copies of a broadly conserved six-residue motif and is predicted to form a three-faced, right-handed β-helix structure. The predicted structure displays a strip of basic residues along each of the three long edges, creating a highly basic surface arranged in a distinct spatial configuration. It is possible that this arrangement facilitates interaction with a partner possessing a complementary, acidic surface. Functionally relevant β-helical structures have been identified in several classes of proteins, including pectate lyase (42) and antifreeze proteins (43,44). Notably, the folding and assembly of many triple β-helices depend on a registration signal that regulates the correct three-dimensional structure formation (45). To date it is unknown whether mitosis signals, such as phosphorylation, also regulate the formation of this putative β-helical structure in ZNF512B.

To explore the possibility of direct tubulin binding, we performed multiple AlphaFold simulations to predict interactions between the IN domain and tubulin; however, no binding was predicted. Given that the spindle apparatus comprises hundreds of different proteins (46,47), it remains challenging to identify the direct binding partner of ZNF512B. In our previously published lf-qMS analysis of GFP-ZNF512B interactors in asynchronously growing cells (21), we identified several potential candidates. These include Cytoskeleton-Associated Protein 2 (CKAP2), which is essential for microtubule dynamics and proper chromosome segregation (48), Growth Arrest Specific 2 Like 3 (GAS2L3), a tubulin- and actin-binding protein involved in cytokinesis and known to interact with the chromosomal passenger complex (49,50), and the Non-SMC Condensin I Complex Subunits G (NCAPG) and H (NCAPH), both implicated in the process of mitosis (51,52). However, AlphaFold simulations did not predict binding of ZNF512B to any of these candidates. Hence, future experiments, such as mass spectrometry analysis of GFP-ZNF512B immunoprecipitates from cells arrested in mitosis, may help to identify its direct interaction partner(s).

Since elevated levels of ZNF512B cause a pronounced delay in metaphase exit, while its depletion accelerates cell proliferation, it is tempting to speculate that ZNF512B plays a direct role in the spindle-dependent orchestration of forces required for sister chromosome segregation. Although our limited microscopic analyses did not reveal major defects in spindle formation, cells appeared to arrest in metaphase, suggesting that mechanisms at the metaphase check point may be affected (53). This could involve alterations in mechanical forces that directly influence microtubule polymerization (54) or depolymerization dynamics (55). Alternatively, ZNF512B may impact regulatory components of the metaphase-to-anaphase transition, the so-called spindle assembly checkpoint (SAC) (56).

Another notable observation is that loss of ZNF512B leads to faster proliferating mESCs that fail to properly differentiate into beating CMs or NPCs. This differentiation defect likely results from a combination of mitotic abnormalities and transcriptional deregulation, as ZNF512B also functions as a transcriptional repressor (14,21). Supporting our findings, a recent preprint from the group of Charles G. Bailey reported that ZNF512B may play a role in neurogenesis, based on RNAi-mediated depletion experiments in NTERA-2 neural cells (57).

ZNF512B is not the first zinc finger protein reported to have multiple functions, including roles as a transcriptional repressor, regulator of differentiation, and spindle microtubule-associated factor. Other examples include Kaiso, a BTB/POZ domain-containing zinc finger repressor that has been implicated in the regulation of cell cycle progression and cell proliferation in cancer (58), and ZNF207 (BuGZ), a C2H2-type zinc finger protein. Different isoforms of ZNF207 are involved in loading spindle assembly checkpoint proteins to kinetochores (59) and have also been identified as key drivers of human ESC differentiation into ectoderm lineage (60).

In summary, our findings underscore that zinc finger proteins are not exclusively nuclear transcription factors but can also act as regulators of mitotic progression as well as of stem cell differentiation.

## Data Availability

Raw and processed RNA-seq files are deposited in GSE303898.

## Supplementary Data

Supplementary Data are available online.

## Acknowledgements

We thank all current and past members of the Hake team for practical help, support and ideas, especially Sonja Sahner for her relentless support in organizing the lab, and Tyra Grundig, Rosalie Habermehl and Elias Haddad for experimental assistance. We thank Prof. Dr. Cristina Cardoso (Technical University of Darmstadt, Germany) for providing us with the pc1116-H2B-mRFP plasmid.

## Author Contributions Statement

LWP, TMW and SBH conceived of this study. LWP performed transfections, IPs, IFs quantitative live-cell imaging (Cytena) and RNA-seq. TMW cloned several GFP-ZNF512B constructs and performed some transfections with the support of FD. ND helped with cell staining and microscopy analysis. LWP and TMW performed immunoblots with the assistance of JoL. JJG bioinformatically analyzed RNA-seq data with the help of LWP. JiL and CK helped with mESC differentiation experiments. SW performed transfections and Spinning-Disk live-cell imaging with the advice of AB. EK performed AlphaFold2 and 3 simulations with the help from JPM. SBH, TMW and LWP wrote the manuscript with support and help from all other co-authors.

