## Supplementary Information and Supplementary Figures for "ZNF512B associates with mitotic spindles, regulates metaphase exit and is crucial for stem cell differentiation"

Running title: ZNF512B regulates metaphase exit and cell differentiation

Keywords: ZNF512B / mitotic spindle / metaphase exit / stem cell differentiation / NuRD

### Supplementary Material and Methods

#### *Antibodies*

| Antibody | Host | Supplier | Order Number | Application | Dilution |
|---|---|---|---|---|---|
| α-CHD4 | Mouse | Abcam | ab70469 | IF | 1:100 |
| α-GFP | Mouse | Roche | 11814460001 | IB | 1:3,000 |
| α-H3 | Rabbit | Abcam | ab1791 | IB | 1:5,000 |
| α-H3S10ph | Mouse | Active Motif | 39636 | IB | 1:1,000 |
| α-RBBP4 | Rabbit | Abcam | ab79416 | IF | 1:100 |
| α-alpha-Tubulin | Mouse | Active Motif | 39527 | IF | 1:100 |
| | | | | IB | 1:3,000 |
| α-ZNF512B | Rabbit | BJ-Diagnostik BioScience GmbH | not applicable | IF | 1:100 |
| α-ZNF512B | Rabbit | Atlas Antibodies | HPA006688 | IB | 1:1,000 |
| anti-Mouse IgG (H+L), HRP | Goat | Invitrogen | 31430 | IB | 1:20,000 |
| anti-Rabbit IgG (H+L), HRP | Goat | Invitrogen | 31460 | IB | 1:20,000 |
| F(ab')2 anti-Rabbit IgG (H+L) Cross-Adsorbed, Alexa Fluor™ 488 | Goat | Invitrogen | A-11070 | IF | 1:200 |
| F(ab')2 anti-Rabbit IgG (H+L) Cross-Adsorbed, Alexa Fluor™ 594 | Goat | Invitrogen | A-11072 | IF | 1:200 |
| F(ab')2 anti-Mouse IgG (H+L) Cross-Adsorbed, Alexa Fluor™ 594 | Goat | Invitrogen | A-11020 | IF | 1:200 |

#### *Primers and Oligos*

| Target | Application | Forward (5' to 3') | Reverse (5' to 3') |
|---|---|---|---|
| ZNF512B | Cloning of GFP-ZNF512B_IN | TCATCGTTCGAAGTCCGGATCTATGGACCGACCCCTGC | TAGTAAGCGGCCGCTCACACCTTCCGCTTCTTCTTGGGTGTGGCACGAGGTGCTTT |
| | Cloning of GFP-ZNF512B_IC | TCATCGTTCGAAGTCCGGATCTGGGAGGAACAGTGGTAAGAAAAGGG | TAGTAAGCGGCCGCTCAGACGGCTTCCCCGC |
| | Cloning of *X. laevis* GFP-Znf512b | CTTCGAAGTCCGGATCTGCTCATCCAATATCTGCTGGGT | CGCGGCCGCTCACTATTACTTCACCCTTCGGGATGG |
| Generation of *Znf512b* mESC KOs | Amplifying genomic region for left homology arm | GGACTCACCTGCTTGAACCTTG | GAGAGCTTGGTTCTCGGCTTT |
| | Amplifying genomic region for right homology arm | ATGGAATCACATGGACCACCC | TAGAGGGACAGGAGGCTTACC |
| | Left homology arm + 12bp pUC18 and | GATTACGAATTCGGCAGGTAGGATAGCTCAG | GCCCTTGCTCACCATCCCTATAGTACAGGAGACAGT |

| | | | |
|---|---|---|---|
| | mCherry overhangs for Gibson Cloning | | |
| | Left homology arm + 12bp pUC18 and Puromycin overhangs for Gibson Cloning | GATTACGAATTCGGCAGGTAGGATAGCTCAG | CTTGTACTCGGTCATCCCTATAGTACAGGAGACAGT |
| | Right homology arm + 12 bp triple terminator and pUC18 overhangs for Gibson Cloning | GATGAATAAATTTCAGCAAGCCCATCGG | AGTGCCAAGCTTGTTTGGCCAGGCCTGT |
| | gRNA 1a/b | CACCGGTATGTCCACTGTGGACCAC | AAACGTGGTCCACAGTGGACATACC |
| | gRNA 1c/d | CACCGGCCCCACTCTGCTTTGACCC | AAACGGGTCAAAGCAGAGTGGGGCC |
| | gRNA 2a/b | CACCGTGGTGACCGGCCGGCTGACT | AAACAGTCAGCCGGCCGGTCACCAC |
| | gRNA 2c/d | CACCGCACCATCAGCCGGCCTGTTG | AAACCAACAGGCCGGCTGATGGTGC |
| | Amplification of pUC18 backbone with homology arm overhangs | GGCCTGGCCAAACAAGCTTGGCACTGGC | ATCCTACCTGCCGAATTCGTAATCATGGTCATAGCT |
| | Amplification of mCherry with homology arm overhangs | ACTATAGGGATGGTGAGCAAGGGCGAG | TGGGCTTGCTGAAATTTATTCATCCCACATAACTGAAATTTTATACC |
| | Amplification of puromycin with homology arm overhangs | ACTATAGGGATGACCGAGTACAAGCCCA | TGGGCTTGCTGAAATTTATTCATCCCACATAACTGAAATTTTATACC |
| Cloning of pIRES-RFP-H2B | Amplification of RFP-H2B | CTACCGGTCGCCACCATGCCAGAGCCAGCGAAG | AGATCTGAGTCCGGATTAGGCGCCGGTGGAG |
| Genomic PCR | Amplification of pIRES | TCCGGACTCAGATCTCGAGC | GGTGGCGACCGGTAGC |
| qPCR Primers | *Znf512b* | AGGAAGTGCGGAAGAAGGTG | TCCGGGCCACCTGGG |
| | *Ctnnb1* | CGTGTCCTGTGAAGCCCG | CCAACTCCATCAGGTCAGCTTG |
| | *Tgfbr3* | CAGACCAATGGCTACTCGGG | GAGCCTGCACCACAATAGAGT |
| | *Dkk3* | AAGGCAAGAGGAGCCATGAATGT | TGGTCTCCACAGCACTCACT |
| | *Wnt3a* | TGGCTCCTCTCGGATACCTCT | CACAGCCAAGGACCACCAGAT |

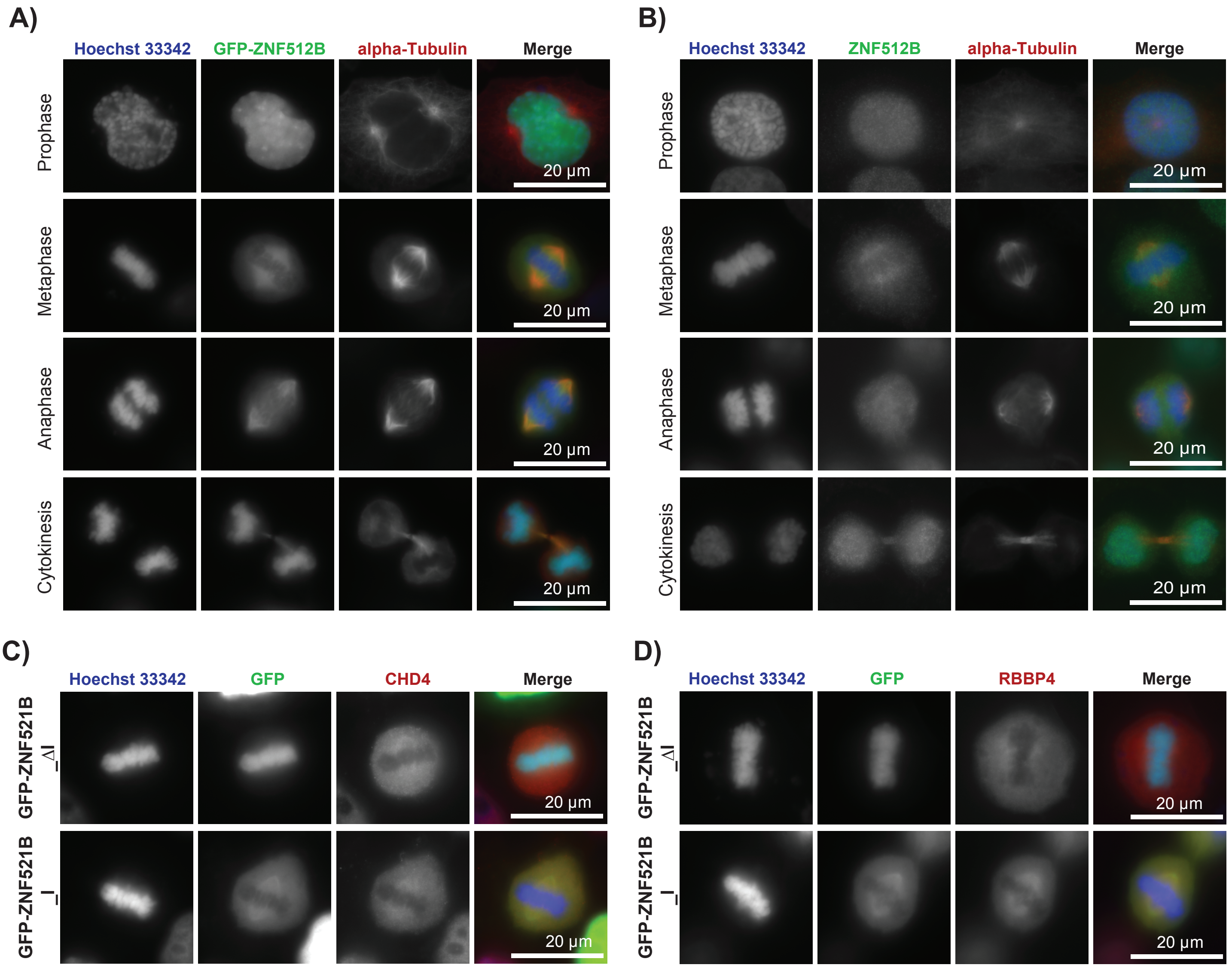


**Supplementary Figure S1:**
**(A)** Immunofluorescence microscopy pictures of HeLaK cells transfected with GFP-ZNF512B (green). DNA (blue) was visualized by Hoechst 3342 and microtubules/mitotic spindles by anti-alpha-tubulin (red) staining of cells in prophase, metaphase, anaphase and cytokinesis. Scale bar = 20 µm.
**(B)** Immunofluorescence microscopy pictures of HeLaK cells stained with anti-ZNF512B antibody (green). DNA (blue) was visualized by Hoechst 3342 and microtubules/mitotic spindles by anti-alpha-tubulin (red) staining of cells in prophase, metaphase, anaphase and cytokinesis. Scale bar = 20 µm.
**(C, D)** Immunofluorescence microscopy pictures of metaphase HeLaK cells expressing GFP-ZNF512B_I or GFP-ZNF512B_ ΔI (green) and stained with Hoechst 3342 solution (DNA, blue) and anti-CHD4 (**C**) or anti-RBBP4 (**D**) (red) antibodies. Scale bar = 20 µm.

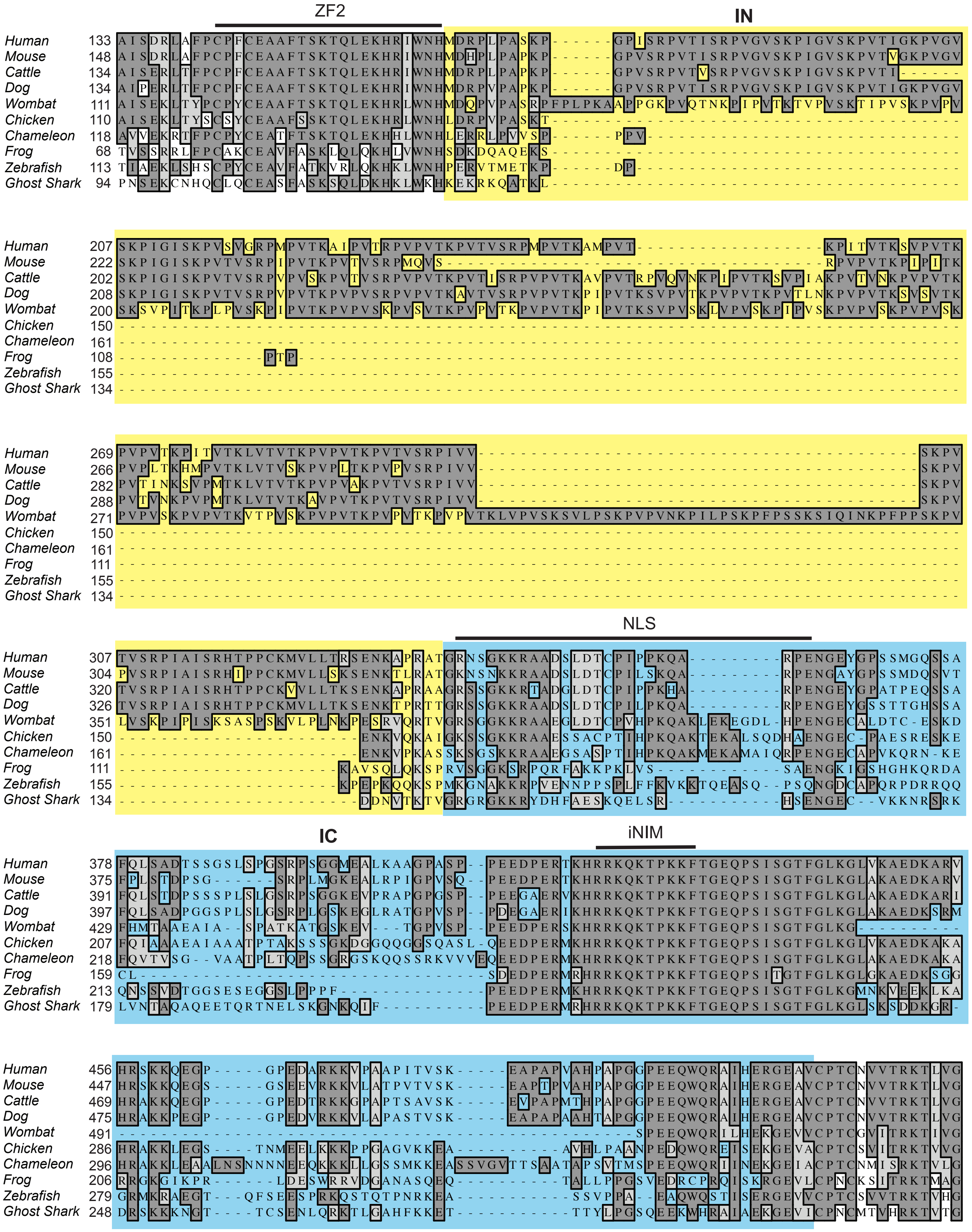

**Supplementary Figure S2: Evolutionary conservation of ZNF512B's internal domain.**
Protein sequences from human (Homo sapiens, Entry Q96KM6), mouse (Mus musculus, Entry Q6ZPW1), cattle (Bos taurus, Entry G3N139), dog (Canis lupus familiaris, Entry A0A8P0NRY1), wombat (Vombatus ursinus, Entry A0A4X2L7X1), chicken (Gallus gallus, Entry A0A8V0ZIQ4), chameleon (Anolis carolinensis, Entry G1KFN9), frog (Xenopus laevis, Entry A0A1L8ELW3), zebrafish (Danio rerio, Entry A0A8M2B8T6) and ghost shark (Callorhinchus milii, Entry A0A4W3IEH9) were obtained from UniProt (uniprot.com). Sequence alignment was performed using MacVector software. Identical amino acids are shaded in dark gray, similar amino acids in light gray and different amino acids in white. Structural features such as zinc finger (ZF), NLS and iNIM regions are indicated by lines. IN is marked by yellow and IC by blue boxes.

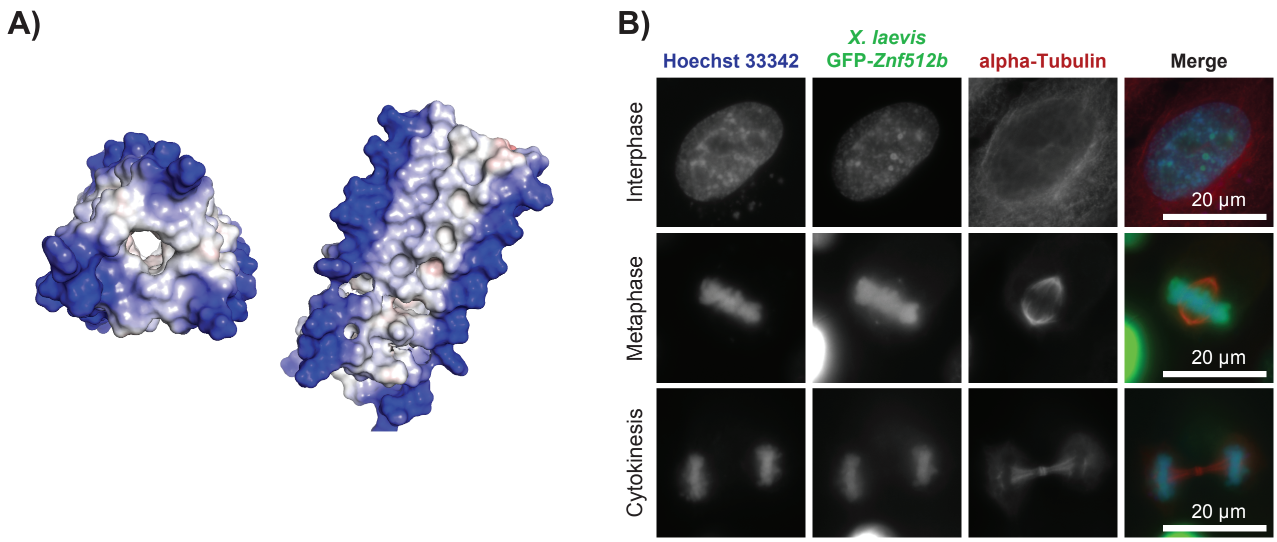


**Supplementary Figure S3: Function and structure of ZNF512B's N-terminal internal domain.**
**(A)** Electrostatic surface prediction (calculated in Pymol 3.1) of the AlphaFold2 prediction for human ZNF512B's IN domain. Blue, gray and red indicate positive, neutral and negative electrostatic potential, respectively. Shown are two views rotated 90°.
**(B)** Immunofluorescence microscopy pictures of HeLaK interphase (top) and metaphase (bottom) cells transfected with GFP (control) and GFP-tagged Xenopus laevis Znf512b constructs (green). DNA (blue) was visualized by Hoechst 3342 and microtubules/mitotic spindles by anti-alpha-tubulin (red) staining. Scale bar = 20 µm.

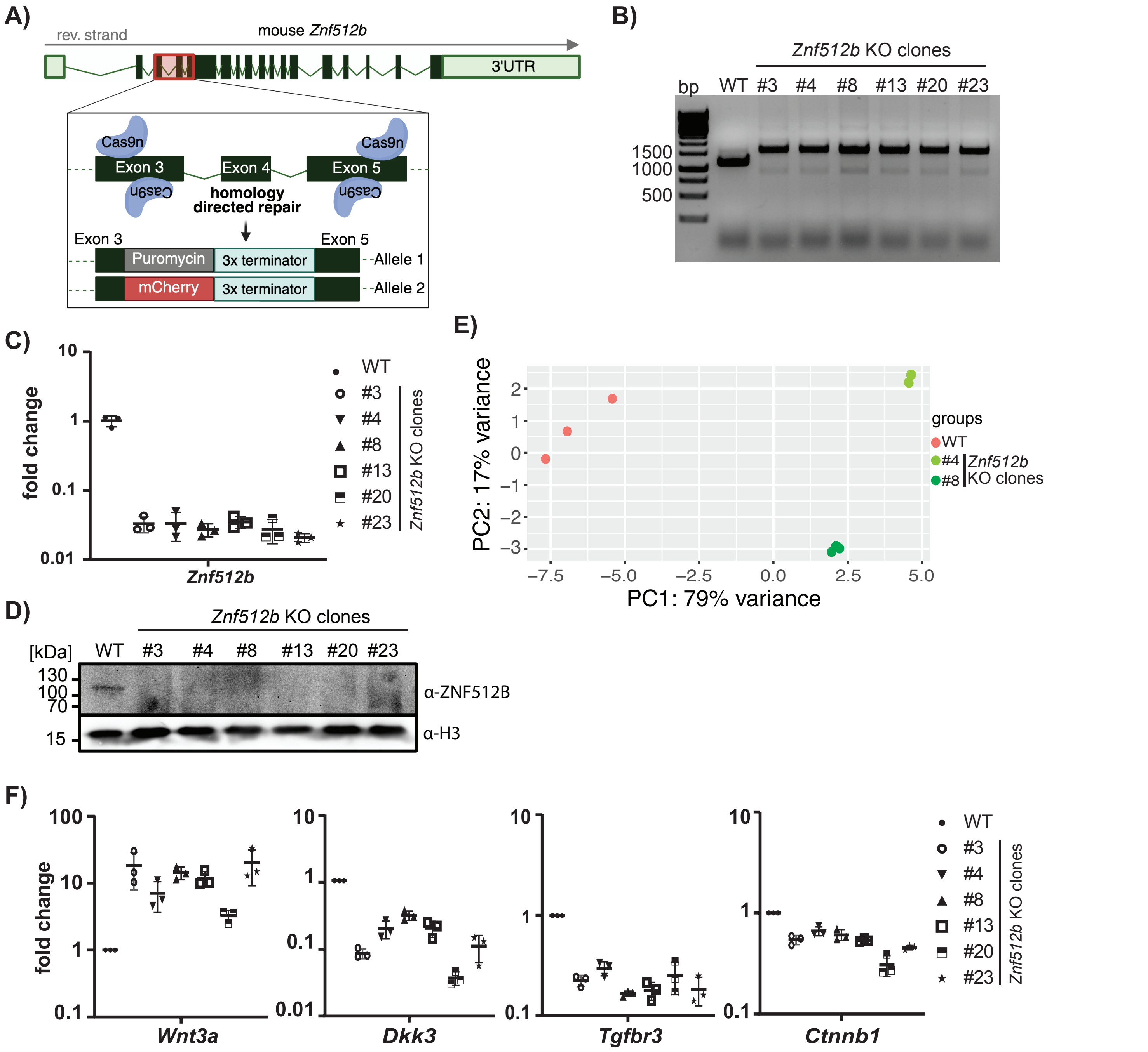


**Supplementary Figure S4: Generation and analysis of Znf512b KO mESCs.**

**(A)** Schematic depicting CRISPR/Cas9-based generation of Znf512b KO mESC clones by introducing triple transcriptional terminator sequences together with either puromycin or mCherry selection markers in between exons 3 and 5 of the murine Znf512b gene.

**(B)** Agarose gel separation of PCR products after PCR amplification of the murine Znf512b locus using genomic DNA isolated from WT and six naïve Znf512b KO mESC clones (#3, #4, #8, #13, #20, #23).

**(C)** RT-qPCR analysis of Znf512b mRNA expression in naïve WT and six Znf512b KO mESC clones (#3, #4, #8, #13, #20, #23) normalized to HPRT expression. Indicated is the fold change in expression compared to WT. Error bars represent the standard deviation.

**(D)** Immunoblots of nuclear extracts derived from naïve WT and six Znf512b KO mESC clones (#3, #4, #8, #13, #20, #23) stained with anti-ZNF512B and anti-H3 (loading control) antibodies.

**(E)** Principle Component Analysis (PCA) of RNA-seq data from three replicates derived from naïve WT (red) and Znf512b KO mESC clones #4 (light green) and #8 (dark green).

**(F)** RT-qPCR analysis of differentially expressed genes Wnt3a, Tgfbr3, Dkk3 and Ctnnb1 in naïve WT and Znf512b KO mESC clones normalized to HPRT expression. Indicated is the fold change in expression compared to WT. Error bars represent the standard deviation.

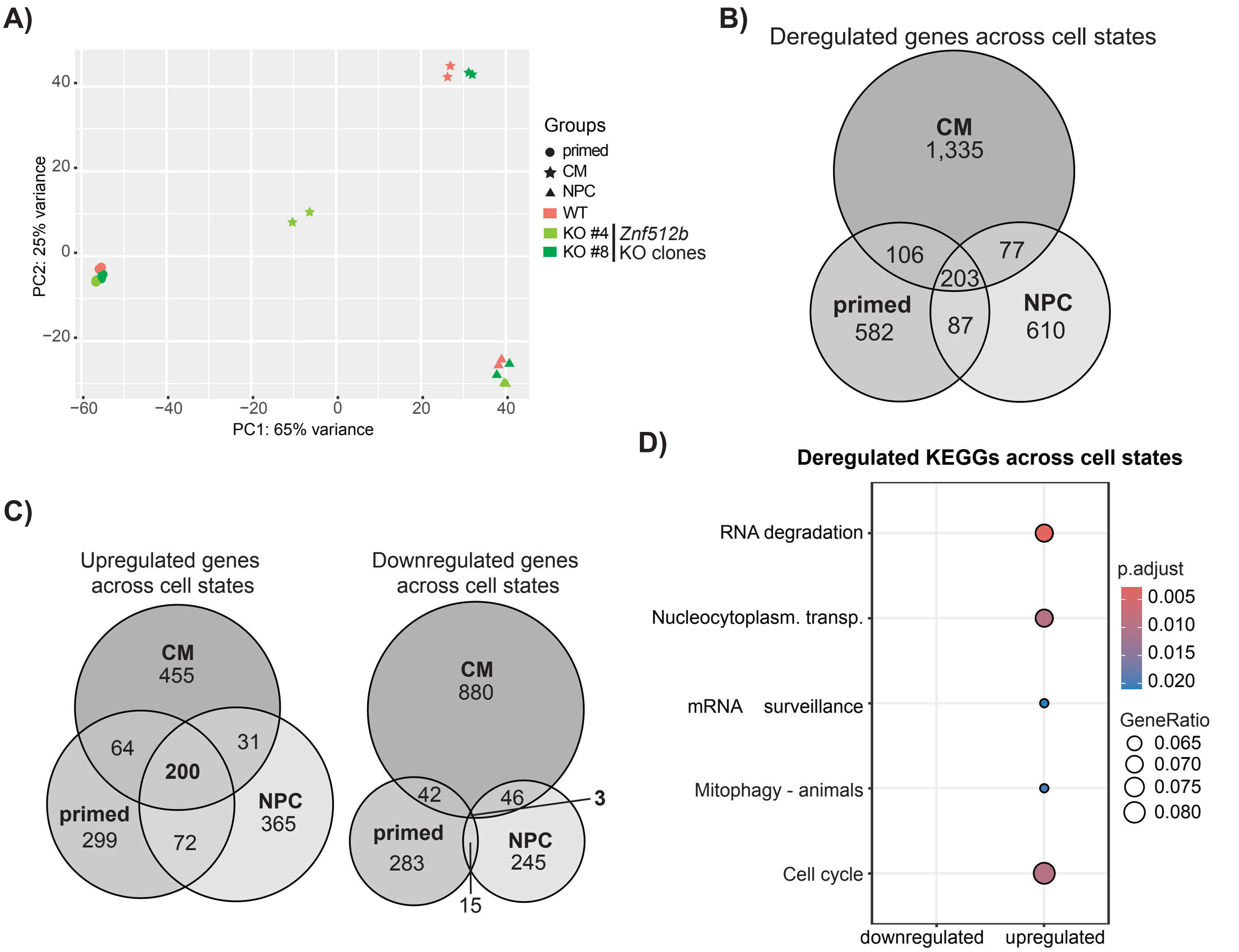


**Supplementary Figure S5: ZNF512B depletion results in upregulation of genes associated with the cell cycle across all three differentiation stages.**

**(A)** Principle Component Analysis (PCA) of RNA-seq data from two replicates derived from WT (red) and Znf512b KO mESC clones #4 (light green) and #8 (dark green) in the primed (pr), NPC or CM stages.

**(B)** Venn diagram of shared deregulated genes in Znf512b KO clones #4 and #8 compared to WT mESCs across all three stages.

**(C)** Venn diagrams of shared downregulated (left) and upregulated (right) genes in Znf512b KO clones #4 and #8 compared to WT mESCs in the primed, NPC and CM stages (see **B**).

**(D)** KEGG pathway analysis of shared deregulated genes in both Znf512b KO clones across all three stages (203 genes shown in **B**).

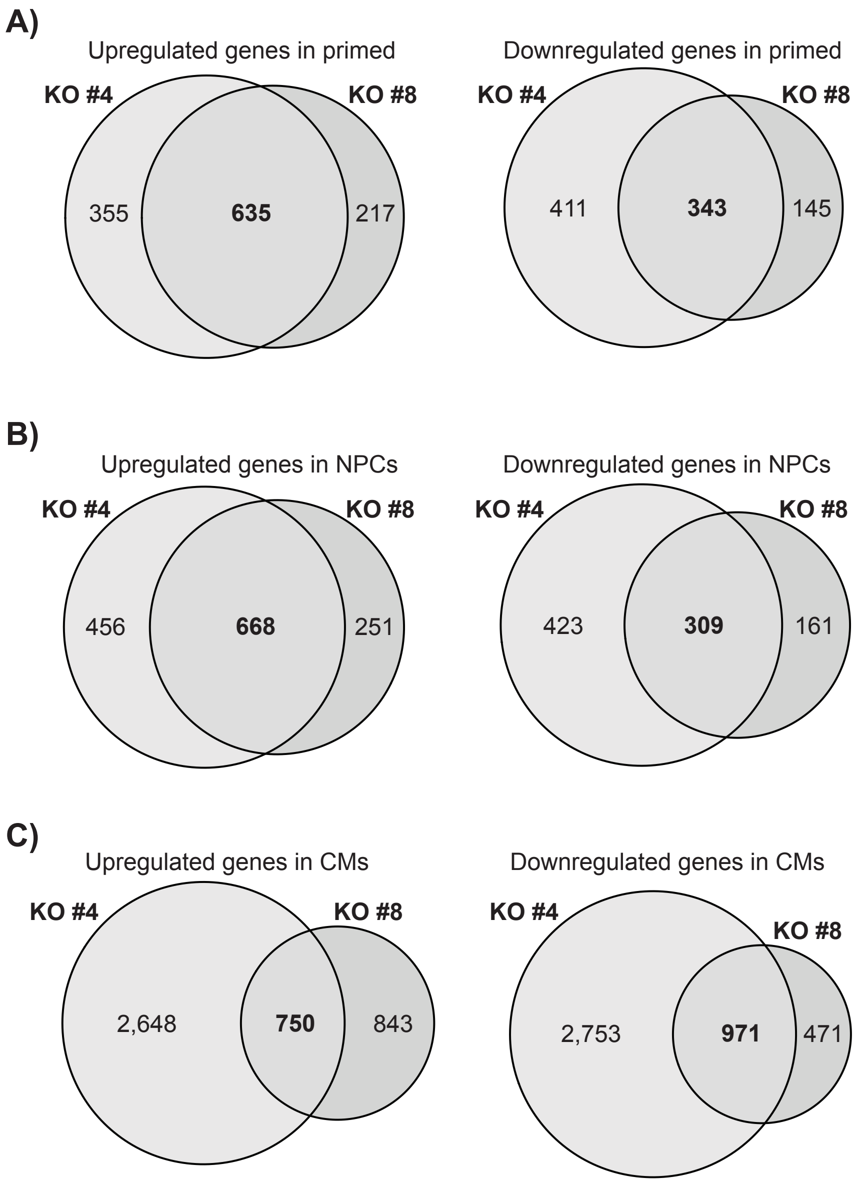


**Supplementary Figure S6: ZNF512B depletion results in deregulated gene expression during mESC differentiation into CMs and NPCs.**
**(A-C)** Venn diagrams of downregulated (left) and upregulated (right) genes in Znf512b KO clones #4 and #8 compared to WT cells in the primed (**A**), NPC (**B**) and CM (**C**) stages.

**Movie Legends:**

**Movie 1:**

Live-cell 3D reconstruction of a HeLaK cell in metaphase transiently expressing RFP-H2B and GFP-ZNF512B. Twenty z-stacks were acquired with a Spinning-Disc microscope at 0.65 µm intervals and reconstructed using Imaris. Scale bar = 5 µm.

**Movies 2–8:**

Videos of live-cell imaging from different time points of HeLaK cells stably expressing RFP-H2B (red) transfected with GFP (Movie 2), GFP-ZNF512B (Movie 3), GFP-ZNF512B_K423A (Movie 4), GFP-ZNF512B_ΔI (Movie 5), GFP-ZNF512B_I (Movie 6), GFP-ZNF512B_IN (Movie 7) and GFP-ZNF512B_IC (Movie 8). Cells were monitored over a 36-hour time period, with images acquired every 20 minutes using the CELLCYTE X (Cytena) live-cell imaging system. Movies are shown with 2.5 frames per second. Scale bar = 100 µm.

**Supplementary Tables Legends:**

**Supplementary Table 1:**

RNA-seq data of shared deregulated genes in naïve *Znf512b* KO clones #4 and #8 compared to naïve WT mESCs.

**Supplementary Table 2:**

RNA-seq data of shared deregulated genes in *Znf512b* KO clones #4 and #8 compared to WT cells in primed, NPC and CM stages.
